# Identification of unusually disulphide-bonded insulin forms using mass spectrometry and thermolysin cleavage

**DOI:** 10.1101/2022.03.11.484026

**Authors:** Dillon Jevon, Kyung-Mee Moon, Leonard J. Foster, James D. Johnson

**Affiliations:** Diabetes Research Group, Life Sciences Institute, Department of Cellular and Physiological Sciences & Department of Surgery, Vancouver, Canada, V6T 1Z3; Charles Perkins Centre, School of Medical Sciences, University of Sydney, Camperdown, Australia, 2006; Department of Biochemistry and Molecular Biology, University of British Columbia, Vancouver, Canada, V6T 1Z3

**Keywords:** diabetes, pancreatic beta-cells, neoepitopes

## Abstract

Insulin is an essential hormone made by the pancreatic beta-cells in the islets of Langerhans. Beta-cells produce more insulin protein than virtually all other cellular proteins combined. Dysfunction in the process of insulin synthesis can lead to disease, including rare forms of monogenic diabetes. Specifically, aberrant intra-insulin and inter-insulin disulphide bonds have been implicated in the pathology of type 1 diabetes and type 2 diabetes, respectively. In type 1 diabetes, misprocessed insulin isoforms may be neoepitopes that kick-start and/or exacerbate the auto-immune response. In type 2 diabetes, aberrant disulphides form insulin dimers that can clog the endoplasmic reticulum and contribute to beta cell dysfunction. To facilitate the study of novel and known insulin neoepitopes and dimers, we present an unbiased and rapid technique for identifying insulin disulphide patterns from pancreatic islet extracts. The basis of this method is the cleavage between insulin’s cysteine residues with the metalloprotease, thermolysin, and subsequent identification of cysteine containing fragments and their partner peptides by LC-MS/MS. Using this technique, we identify 6 aberrant disulphide bonded insulin species, including a previously described type 1 diabetes neoepitope, as well as inter-chain disulphide bonded insulin dimers. Furthermore, using the endoplasmic stress inducer, thapsigargin, we observe increased disulphide errors in a patient donor sample. This approach lays foundations to identify the scope and cause of aberrant insulin disulphide formation in health and disease.

## INTRODUCTION

Insulin’s structure was studied extensively by Frederick Sanger in the mid 20^th^ century, a pursuit which ultimately earned him his first Nobel prize in 1958 (1–7). To determine the disulphide bonding pattern of insulin, Sanger and colleagues employed an acid hydrolysis method in which the peptide bonds of insulin were degraded, while the disulphide bonds were left intact (8). Hydrolytic insulin products were then separated by paper ionophoresis, oxidised, and separated again to determine the amino acid composition of each disulphide bonded peptide fragment. Since then, Edman degradation and mass spectrometry have been used to cleave insulin between its cysteines and resolve the structure of the resultant products (9,10). However, these techniques require the lengthy and specialized preparation of multiple samples to determine insulin’s disulphide bonds. A modern and rapid technique, able to identify all insulin disulphides present in a single sample, is needed to help determine the scope and severity of disulphide mis-formation in type 1 diabetes and type 2 diabetes.

Disruption of proinsulin disulphide bonding and folding is associated with several disease states, such as maturity-induced diabetes of the young and, more recently, T1D. Irregular insulin-derived peptides have been recognised by autoimmune CD4^+^ and CD8^+^ T cell clones from donors with type 1 diabetes (T1D), but not control donors (11). In one study, an irregular insulin fragment was characterized by an intra-chain disulphide bond between cysteine residues 6 and 7 of the insulin A chain (A6:A7), which normally form disulphides with insulin residues A11 and B7, respectively (11). Donor CD4^+^ T cell clones from T1D donors, but not controls, recognized the epitope formed by this insulin fragment. While biological conditions that might cause erroneous disulphide bonding to occur in T1D pathology have not been identified, endoplasmic reticulum (ER) stress has been the subject of recent study in the field of T1D. Insulin disulphides form in the ER, and ER stress precedes occurrence of insulitis in the non-obese diabetic mouse model of T1D. Furthermore, the ER stress inducer, thapsigargin, has been used to induce production of an alternative reading frame insulin mRNA product which like the A6:A7 bonded insulin fragment above is recognised by T cell clones from human T1D donors and not controls (12–15). These peptides have become known as “neoepitopes” from the hypothesis that new (neo) epitopes form from erroneous protein production and become antigens which initiate or exacerbate the auto-immune attack (16). Furthermore, the detection of insulin polymers in type 2 diabetes (T2D) suggest inter-molecular insulin disulphide bonds are also pathologically significant in this disease (17). To date, the presence of erroneous insulin disulphides has been observed by directed study to specific insulin fragments suspected of harbouring an aberrant disulphide bond. The discovery of pathologically relevant irregular insulin disulphide warrants development of an unbiased technique that can identify insulin disulphide bonding patterns.

Thermolysin is a thermostable metalloprotease which preferentially cleaves on the N terminus of hydrophobic and bulky amino acid residues (18). Conveniently, insulin contains at least one of these residues present between each cysteine residue, except the adjacent A-chain cysteines, A6:A7. Digestion of non-reduced insulin with thermolysin produces peptide fragments which are bound to partner fragments through single disulphide bonds. Using tandem mass spectrometry, it is possible to identify all peptide fragments and their binding partners, determining the disulphide patterns by identifying the partner fragment bound to each cysteine residue. In the case of a disulphide bond between cysteine residues A6:A7, an intra-chain disulphide can be detected by a −2.016 Dalton shift corresponding to the loss of two hydrogen atoms (9). Here, we present a mass spectrometry protocol which uses thermolysin digestion and omits disulphide reduction to effectively detect six insulin disulphide conformations present from acid-ethanol extracts of human pancreatic islets.

## METHODS

### Primary Human Islet Culture and Insulin Extraction

Human cadaveric islets from donors (R305, R310, R326) were obtained from the University of Alberta Islet Core (donor information can be found at the following website - https://www.epicore.ualberta.ca/isletcore/Default) and arrived on mornings after overnight shipping at ambient temperature. Islets (2500 IEQ) were resuspended in 5.5mM glucose RPMI (Sigma–Aldrich, St Louis, MO) supplemented with 10% (v/v) FBS (Life Technologies, Burlington, ON) and penicillin/streptomycin (100 μg/ml; Life Technologies), and cultured overnight. After overnight culture, islets were centrifuged at 200 x g for 10 min and the media removed. The pellet was resuspended in 200 μl acid ethanol extraction buffer (1.5% HCl, 25% acetic acid, 70% ethanol). The suspension was transferred to 1.5mL sample tubes and vortexed on high for one minute to extract insulin. The sample tubes were then centrifuged at 27,200 x g for 10 minutes and the soluble fraction collected.

### Mass Spectrometry Sample Preparation

Acid-ethanol extract was evaporated by vacuum centrifugation and the protein reconstituted in 100ul of thermolysin digestion buffer (50mM Tris, 0.5mM CaCl_2_, pH 8.0), with protein concentration then measured by nanodrop spectrophotometry. Sample pH was acidic after reconstitution in digestion buffer and ammonium bicarbonate (Fisher, USA) was added until pH6.0 before digestion. Thermolysin (Promega; Madison, WI) was added to sample in a 1:20 enzyme:protein ratio and the digestion proceeded at 90°C for 3.5 h. Standard reduction procedures used in proteomic preparations were omitted and thermolysin’s high digestion temperature substituted for commonly used chemical denaturants. After digestion, samples were cleaned and eluted by stop-and-go extraction as described by Rappsilber and colleagues (19).

### Mass Spectrometry Acquisition Conditions and Data Analysis

We employed a discovery proteomics approach to map proinsulin-related peptides from human β cells, including incorrectly disulphide-bonded peptides (20). Insulin peptides were identified by a quadrupole time of flight mass spectrometer (Impact ii; Bruker Daltonics) coupled to an easy nano lc 1000 hplc (Thermo Fisher) as described (21). Because the insulin peptides required thermolysin protease, singly charged peptides were also fragmented. Spectra were searched against Uniprot’s reviewed homo sapiens reference database (up000005640: 20,384 entries, downloaded on September 8, 2021) with additional custom insulin sequence variations and the software’s built-in common contaminants using both maxquant version 1.6.17.0 (22) and byonic version 4.0.12 (23). In both software packages, the enzyme cleavages were set to cut at n-terminal ends of lviafm amino acid residues, with maximum missed cleavages of 3. The peptide mass tolerance was set to 15 ppm in both softwares, and fragment ion mass tolerance was set to 30 ppm and 40 ppm for byonic and maxquant, respectively. In both searches, variable modification of oxidation was set, for hmw residues for byonic and only m residue for maxquant. Variable modification of protein acetylation was added to maxquant search parameters. To detect the disulphide bonding patterns on ins peptides, disulfide bond search option was enabled on byonic with disulfide protein groups set as 1, 2, 3, where the first 3 entries in the fasta file were the three main insulin chains. To retrieve the extracted ion chromatogram value (semi-quantitative) of the crosslinked peptides identified in byonic, the corresponding uncalibrated m/z species intensity value was retrieved from maxquants allpeptides. Txt output, with 5 ppm mass difference and 30 seconds of retention time.

## RESULTS

### Validation and characterization of the extraction protocol

The goal of the present study was to characterize insulin protein isoforms (Fig. 1) from human pancreatic islets. However, first we needed to assessed the purity of our extraction protocols in order to generate samples enriched for insulin. Acid-ethanol extraction from human donor islets was found to produce samples wherein insulin was in high abundance relative to contaminating peptides (Fig 2a). Furthermore, human proinsulin peptide coverage reached 100% in all donor islet samples. Acid-ethanol extraction provided samples which could be quickly processed without the need for further purification and our results likely reflect optimal insulin solubility in an acid-ethanol solution. Acid-ethanol extraction has been standard method for insulin isolation for decades (24,25).

**Figure 1.**
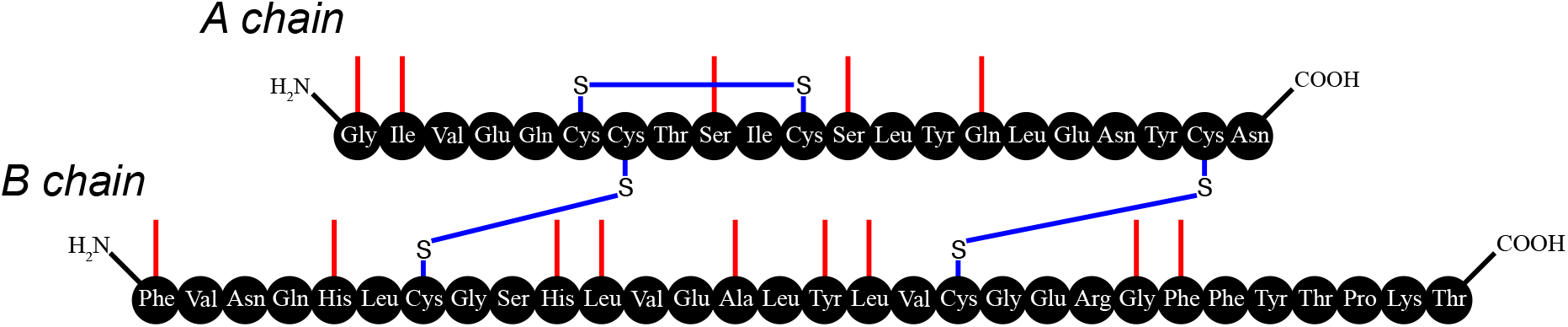
Thermolysin digestion sites in human insulin. Insulin schematic showing thermolysin cleavage sites (provided by Expasy PeptideCutter). Cleavage sites are shown in red (cleavage to C-terminal side of residue). Insulin’s native disulphides are shown in blue.

**Figure 2.**
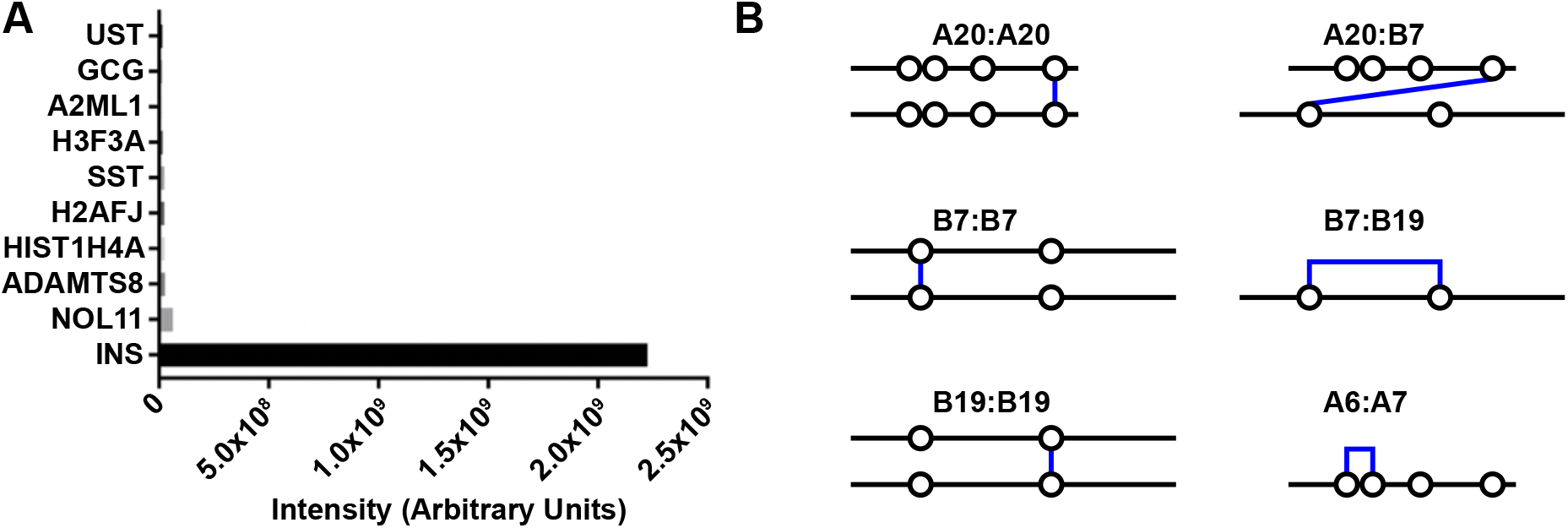
Insulin abundance and aberrant disulphides from human islet acid-ethanol extracts. **A)** Ion intensity corresponding to the 10 most abundant proteins observed by mass spectrometry in acid-ethanol extracts of human donor islets. **B)** Summary of all aberrant disulphides detected in acid ethanol extracts of human islets. Hollow circles correspond to cysteine residues. Blue lines, disulphide bonds.

### Identification of thermolysin cleavage products

Thermolysin cleaves on the N-terminal side of hydrophobic or bulky amino acid residues (26–28), which are present between insulin’s cysteine residues (Fig 1). By digesting non-reduced samples, otherwise separate cleavage products remained linked by disulphide bonds. Insulin products of interest contained two thermolytic fragments, each containing a cysteine and joined by a disulphide bond. Tandem mass spectrometry of digested insulin identified each peptide and determined the mass of cysteine residue additions, which corresponded to the mass of cysteine containing partner peptides (Fig 2b). In the case of the adjacent cysteines, A6:A7, a mass shift of −2.016 Da corresponded to the loss of two hydrogen atoms due to the presence of an intra-chain disulphide was observed (Fig 3). We found no evidence of peptides that were present in one of the untreated or thermolysin treated sample, but not the other.

**Figure 3.**
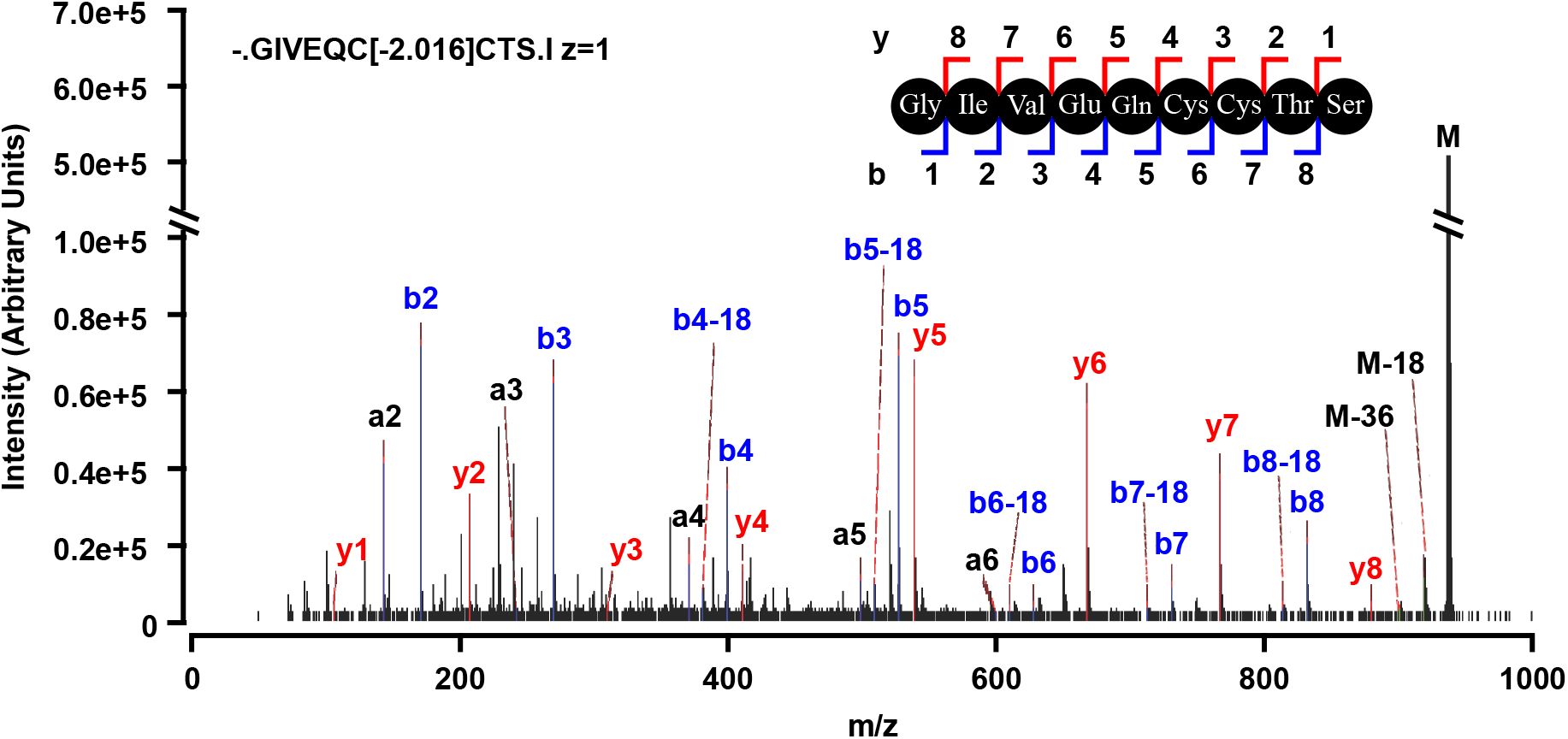
Mass spectra of internally bonded GIVEQCCTS A chain peptide. The intra-chain A6:A7 bonded insulin A chain fragment was detected by a −2.016 dalton mass shift. Red = y ions, blue = b ions.

### Semi-quantification of cleave product abundance

We next performed semi-quantitative analysis to determine if thermolysin treatment might induce a quantitative shift towards specific peptides. This revealed the quantity of erroneous disulphide bonded insulin products to be low relative to canonical insulin disulphide fragments (Fig 4). In two of three samples, there was little to no quantitative difference in peptide abundance between untreated and thermolysin-treated islets. However, one sample had high ion intensity corresponding to the A6:A7 intra-disulphide bonded peptide after thermolysin treatment and not in the donor matched control.

**Figure 4.**
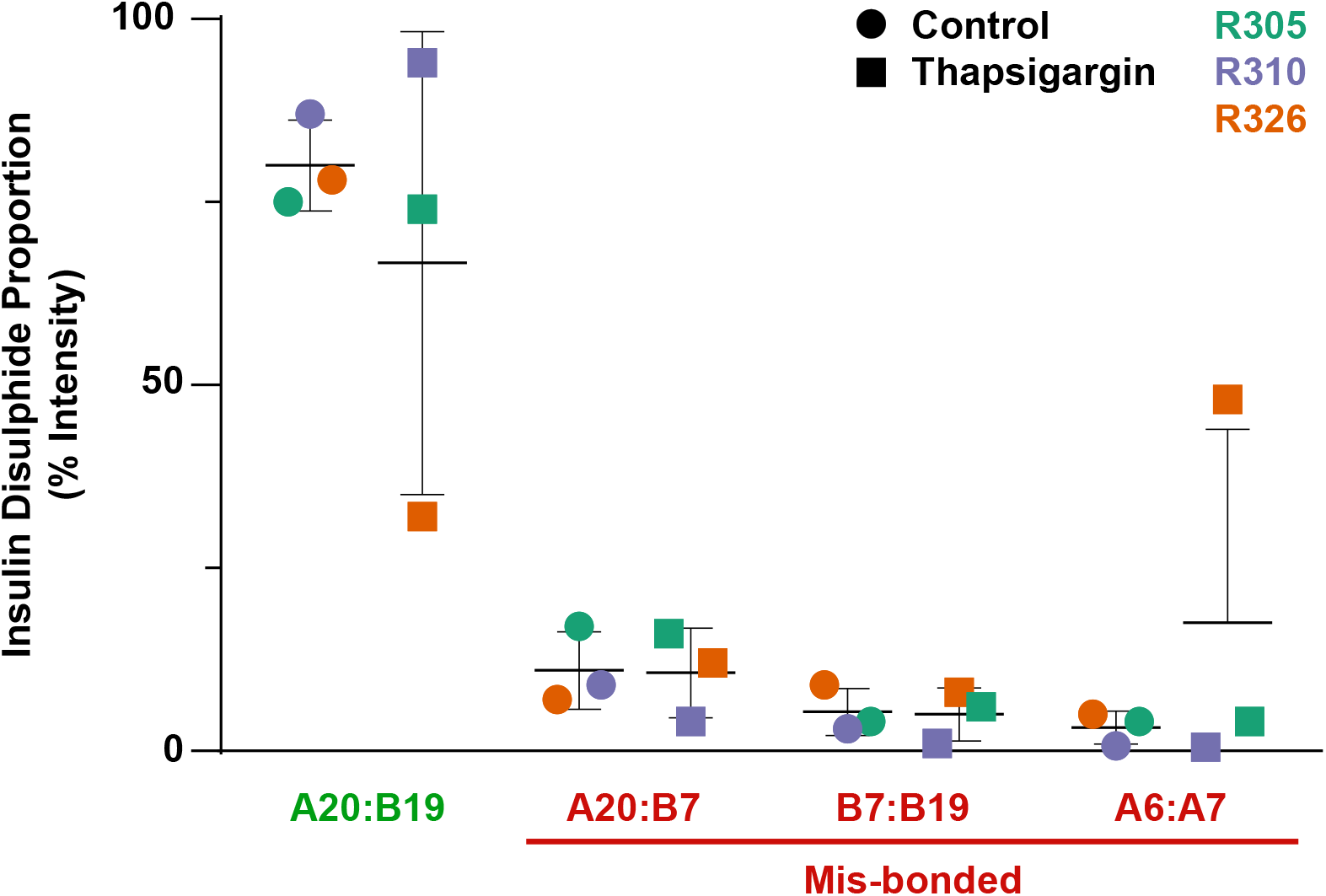
Relative Quantitation of Insulin Disulphide Fragments. Insulin disulphide fragments proportions were quantified relative to the total ion intensity of all insulin disulphide fragments detected in Maxquant seach. Disulphide bonds are annoted based on A or B chain cysteine residue position and colour coded for native or erroneous disulphide. Matched donor islet samples are colour coded. Green=R305, Purple=R310, Orange=R326.

## Discussion

Acid-ethanol extracts of insulin from human islets provided insulin with a diverse array of disulphide bonds needed to validate the above method’s ability to identify a catalogue of disulphide isomers. While the identification of six aberrant disulphide conformations of insulin may seem surprising, the islets used in this study were subject to extraction from cadaveric donors, isolation and overnight shipment at ambient temperature. These non-physiological stressors may be sufficient to cause erroneous disulphide formation. Furthermore, Ryle & Sanger noted disulphide interchange reactions occurred in acidic conditions, albeit at higher temperatures and over longer time periods than were used in this protocol (29). While disulphide interchange may have benefitted this investigation by providing many disulphides for analysis, application of this protocol to study biological (mis)formation of insulin disulphides would benefit from blockage of interchange reactions. Addition of 100 μm thioglycolic acid prevented disulphide interchange in 50% acetic acid over 5 days at 35°C (29) and could be added to acid-ethanol extraction buffers – which in this protocol (excluding time in the vacuum centrifuge) requires only 2 minutes incubation at room temperature.

The relative abundance of disulphide fragments between control and thermolysin treated samples has been compared and relative abundance was inferred from total ion intensity (Fig 4). While there was no significant difference between groups, one sample had increased aberrant disulphide bonding after thapsigargin treatment compared to its matched control (Fig 4, orange). This may indicate that genetic or environmental predisposition could contribute to insulin misprocessing under ER stress. Interestingly, the disulphide produced was a previously described T1D neoepitope (11). Within controls and all but one treatment sample, canonical disulphide bonding constituted a majority of the total ion intensity from thermolytic insulin fragments (Fig 4). We conclude that erroneous insulin disulphides are present in a small amount through virtue of normal biological process or stress through islet isolation and ex-vivo culture and that insulin disulphide errors may occur more frequently under ER stress.

Islets treated with coxsackievirus (30–32) and other ER stressors (33,34) such as tunicamycin could determine whether neoepitope production is increased with physiological stressors associated with T1D when paired with our method. ER stress is a likely cause of insulin disulphide errors as insulin disulphides form in the ER (34,35). Furthermore, insulin production itself contributes to a baseline level of ER stress in beta cells (36,37) and coxsackievirus infection causes ER stress in pancreatic beta cells (35). Including stressors such as coxsackievirus treatment of beta cells in experiments using this thermolysin mass spectrometry protocol could help identify the physiological stressors of neoepitope production. To date, erroneous insulin disulphides (11), defective ribosomal insulin gene products (12) and insulin-GAD41 peptide fusion errors (15) have been identified as immune-provoking molecules in T1D. What causes the production of these erroneously-bonded peptides remains unknown but may yield profound insight into the aetiology of T1D. It is also unclear whether these peptides are secreted from human beta-cells, in either a constitutive or regulated manner, or whether they can bind to and/or activate the insulin receptor. Interestingly, recent studies have explored insulin peptide dimerization to modulate insulin action therapeutically (38).

Our work has introduced a method which allows unbiased classification of insulin neoepitopes formed from disulphide errors and may be paired with drugs and/or stressors to elucidate their root cause. This study opens new avenues for understanding insulin production, beta-cell stress, and diabetes pathogenesis.

## ACKNOWLEDGEMENTS

We thank Pat MacDonald and the University of Alberta Islet Core for the provision of human islets, as well as the islet donors and their families.

